# Predicting Executive Functioning from Brain Networks: Modality Specificity and Age Effects

**DOI:** 10.1101/2023.06.29.547036

**Authors:** Marisa K. Heckner, Edna C. Cieslik, Lya K. Paas Oliveros, Simon B. Eickhoff, Kaustubh R. Patil, Robert Langner

## Abstract

Healthy aging is associated with structural and functional network changes in the brain, which have been linked to deterioration in executive functioning (EF), while their neural implementation at the individual level remains unclear. As the biomarker potential of individual resting-state functional connectivity (RSFC) patterns has been questioned, we investigated to what degree individual EF abilities can be predicted from gray-matter volume (GMV), regional homogeneity, fractional amplitude of low-frequency fluctuations (fALFF), and RSFC within EF-related, perceptuo-motor, and whole-brain networks in young and old adults. We examined whether differences in out-of-sample prediction accuracy were modality-specific and depended on age or task-demand levels. Both uni- and multivariate analysis frameworks revealed overall low prediction accuracies and moderate to weak brain–behavior associations (R^2^ < .07, *r* < .28), further challenging the idea of finding meaningful markers for individual EF performance with the metrics used. Regional GMV, well linked to overall atrophy, carried the strongest information about individual EF differences in older adults, whereas fALFF, measuring functional variability, did so for younger adults. Our study calls for future research analyzing more global properties of the brain, different task-states and applying adaptive behavioral testing to result in sensitive predictors for young and older adults, respectively.

## Introduction

Healthy, cognitive aging is associated with significant structural changes in the brain like cortical thinning, volumetric shrinkage, and decline in white-matter integrity (Park and Reuter-Lorenz, 2008; Reuter-Lorenz and Park, 2014), as well as changes in the functional network architecture (Spreng and Turner, 2019). These brain changes are thought to be accompanied by a decline in cognitive capacities, in which information processing in several cognitive tasks becomes less efficient, especially in demanding tasks that tap into executive functioning (EF) (Park et al., 2002; Park and Reuter-Lorenz, 2008). In contrast, performance appears to remain rather stable in tasks taxing semantic abilities (Salthouse, 1996; Park et al., 2002), implicit memory, or general knowledge (Park et al., 2002). Therefore, the effects of aging on the brain, but also implications on the behavioral performance, are quite heterogeneous. Overall, previous studies indicate that the cognitive system is highly adaptive and dynamic (Greenwood, 2007; Park and Reuter-Lorenz, 2008), which implies that modifications of the neural architecture such as functional reorganization occur to maintain sufficient levels of cognitive functioning.

EF abilities are relevant for goal-directed thought and adaptive behavior in complex environments and are thus critical for everyday life. Rather than being defined as a single process, EF is a multidimensional construct that involves diverse cognitive abilities. Different lines of research suggest three core subcomponents: inhibitory control, working memory, and cognitive flexibility (Lehto, 1996; Miyake et al., 2000; Alvarez and Emory, 2006; Diamond, 2013). Inhibitory control has been linked to controlling one’s attention, thoughts or emotions to attain higher-order or long-term goals. Working memory is associated with holding content in mind and working with it. For instance, when incorporating new information in plans or considering alternatives. Finally, cognitive flexibility is important in the context of changing one’s perspective or adapting to changing rules/demands. At the neural level, EF has been linked to a distributed set of brain regions that have been unified into the so-called multiple-demand network [intraparietal sulcus, inferior frontal sulcus, dorsolateral prefrontal cortex, anterior insula/frontal operculum, pre-supplementary motor area, anterior cingulate cortex], but also to other brain areas, depending on specific task demands (Teuber, 1972; Duncan and Owen, 2000; Duncan, 2010; Miyake and Friedman, 2012; Camilleri et al., 2018).

Earlier studies found age-related differences in EF performance to be partially accounted for by changes in resting-state functional connectivity (RSFC) within brain networks associated with EF (Steffener et al., 2009; Langner et al., 2015; Hausman et al., 2020) and were able to predict EF abilities of previously unseen individuals from RSFC (Reineberg et al., 2015; He et al., 2021). However, in our companion study (Heckner et al., 2023), investigating the same dataset and applying the same data analysis strategy as in the current study but focusing on network specificity, we demonstrated overall low prediction accuracies (as indicated by the root mean squared error [RMSE], mean absolute error [MAE], and correlation coefficient [Pearson’s *r*]) for individual EF performance levels from within-network RSFC. Furthermore, we did not identify any network specificity, that is, EF performance was not better predicted from an EF-related brain network than from EF-unrelated networks (i.e., perceptuo-motor network, whole brain approach). The overall low prediction accuracies and brain–behavior associations (coefficients of determination R^2^ ≤ .04) raised the question of whether the associations found are indeed meaningful. Together with previous research (Finn, 2021; Finn and Bandettini, 2021), these findings challenged the notion that biomarkers for individual EF performance can be found using RSFC patterns.

Since the effects of cognitive aging as well as the behavioral consequences appear quite heterogeneous, MRI metrics capturing different aspects of brain structure and function may need to be applied. A commonly used metric derived from resting-state fMRI, regional homogeneity (ReHo), has shown to be sensitive in identifying age differences during rest (Wu et al., 2007) and offered a better prediction accuracy of crystallized intelligence compared to RSFC (Larabi et al., 2021). ReHo measures the local similarity of a voxel’s time series to its neighboring voxels and is based on the assumption that meaningful brain activity is represented in clusters of neighboring voxels rather than single voxels (Zang et al., 2004). It has been discussed as local connectivity that is necessary to induce global connectivity (Jiang and Zuo, 2016). Another metric derived from resting-state fMRI is fractional amplitude of low-frequency fluctuations (fALFF), which reflects the relative contribution of low-frequency fluctuations within a specific frequency band to the whole frequency range (Zou et al., 2008) and can thus be taken as a measure of functional within-subject brain variability. Previous studies have identified a negative association between functional brain variability, as measured through fALFF, and age. These changes were associated with cortical atrophy, measured through cortical thickness or gray-matter volume (GMV), and a decline in inhibitory control (Hu et al., 2014; Vieira et al., 2020).

As our recent companion paper questioned RSFC’s potential as a biomarker for individual EF abilities, the aim of the current study was to investigate further functional and structural brain metrics for their potential as a biomarker. For this purpose, we defined an EF network (EFN) in the brain by integrating the results of previous neuroimaging meta-analyses (Rottschy et al., 2012; Langner et al., 2018; Worringer et al., 2019), each encompassing diverse facets of EF. As EF-unrelated control networks, we further included a perceptuo-motor (Heckner et al., 2021) and a whole-brain network (Power et al., 2011) for prediction. Then, we examined to what degree individual abilities in three main EF subcomponents (i.e., inhibitory control, cognitive flexibility, and working memory) could be predicted from GMV, RSFC, ReHo, and fALFF within these networks in young and older adults, respectively. For each to-be-predicted EF performance score, we separately sought to predict performance in a high-demand task (representative of increased EF demand) and a low-demand EF control condition. We implemented a linear approach using partial least squares regression (PLSR) for prediction, as previous studies revealed comparable prediction accuracies using a non-linear approach (random forest) or connectome-based predictive modelling (Finn et al., 2015; Shen et al., 2017; Heckner et al., 2023). Overall, we investigated (i) whether one of the structural or functional metrics (GMV, RSFC, ReHo, fALFF) outperforms the others in predicting EF, (ii) if this pattern changes depending on the network, task-demand level, or age group, and (iii) if young and older adults differ in their predictability depending on the structural or functional metric, network, or task-demand level.

## Methods

### Sample

Whole-brain magnetic resonance images of 116 healthy young (age range = 20-40 years, mean age = 26.67, SD = 5.80, 64 females) and 111 old (age range = 60-80 years, mean age = 68.19, SD = 5.66, 72 females) adults were obtained from the publicly available enhanced Nathan Kline Institute - Rockland Sample (eNKI-RS; Nooner et al., 2012). These age bins were chosen to maximize the age variance for studying age-related differences in the association between brain features and behavioral target variables. We excluded participants with acute and/or severe psychiatric or neurological disorders in the past or when currently taking medication presumably affecting brain activity. The re-analysis of the data was approved by the local ethics committee of the Medical Faculty at the Heinrich Heine University Düsseldorf. All participants underwent the same protocol. The sample used was the same as in our companion paper (Heckner et al., 2023). The specific sample used is available upon request.

### Neuroimaging Data Acquisition and Processing

Brain images were acquired on a Siemens TimTrio 3T MRI scanner (Siemens Medical Systems, Erlangen, Germany). T1-weighted structural images were obtained using a MPRAGE sequence [TR = 1.9 s, TE = 2.52 ms, flip angle = 8°, in-plane resolution = 1.0 × 1.0 × 1.0 mm^3^] and further analyzed using SPM12 (Wellcome Trust Centre Neuroimaging, London, https://www.fil.ion.ucl.ac.uk/spm/) and the CAT12 toolbox (Gaser et al., 2022). Whole-brain resting-state fMRI data was obtained using BOLD (blood oxygen level–dependent) contrast [gradient-echo EPI (echo planar imaging) pulse sequence, TR = 1.4 s, TE = 30 ms, flip angle = 65°, voxel size = 2.0 × 2.0 × 2.0 mm^3^, 64 slices, 404 volumes]. Participants were instructed to keep their eyes open and maintain fixation on a central dot. Physiological and movement artifacts were removed from RS data by using FIX (FMRIB’s ICA-based Xnoiseifier, version 1.061 as implemented in FSL 5.0.9; Griffanti et al., 2014; Salimi-Khorshidi et al., 2014), which decomposes the data into independent components and identifies noise components using a large number of distinct spatial and temporal features via pattern classification. Unique variance related to the identified artifactual components is then regressed from the data. Data was further preprocessed using SPM12 and in-house MATLAB scripts. After removing the first four functional dummy volumes, the remaining EPI volumes were corrected for head movement by a two-pass affine registration procedure. First, images were aligned to the initial volume and, subsequently, to the mean of all volumes. The mean EPI image was then co-registered to the gray-matter probability map provided by SPM12 using normalized mutual information and keeping all EPI volumes aligned. Next, the mean EPI image of each participant was spatially normalized to MNI-152 space using the “unified segmentation” approach (Ashburner and Friston, 2000). The resulting deformation parameters were then applied to all other EPI volumes.

### Brain Networks

We used three different networks for our prediction analyses, which were the same as in our companion study (Heckner et al., 2023): (1) An EF-related network (EFN) based on the maximum conjunction of three pertinent meta-analyses investigating working memory (Rottschy et al., 2012), cognitive action regulation (Langner et al., 2018), and multi-tasking (Worringer et al., 2019). The resulting network comprised 50 nodes (i.e., brain coordinates). (2) An EF-unrelated, perceptuo-motor network that integrated visual, auditory, and motor processes and comprised 59 nodes (Heckner et al., 2021). (3) A whole-brain network as a control. We employed Power et al.’s (2011) graph of putative functional areas, which includes 264 nodes. All networks are displayed in Figure 1.

**Figure 1.**
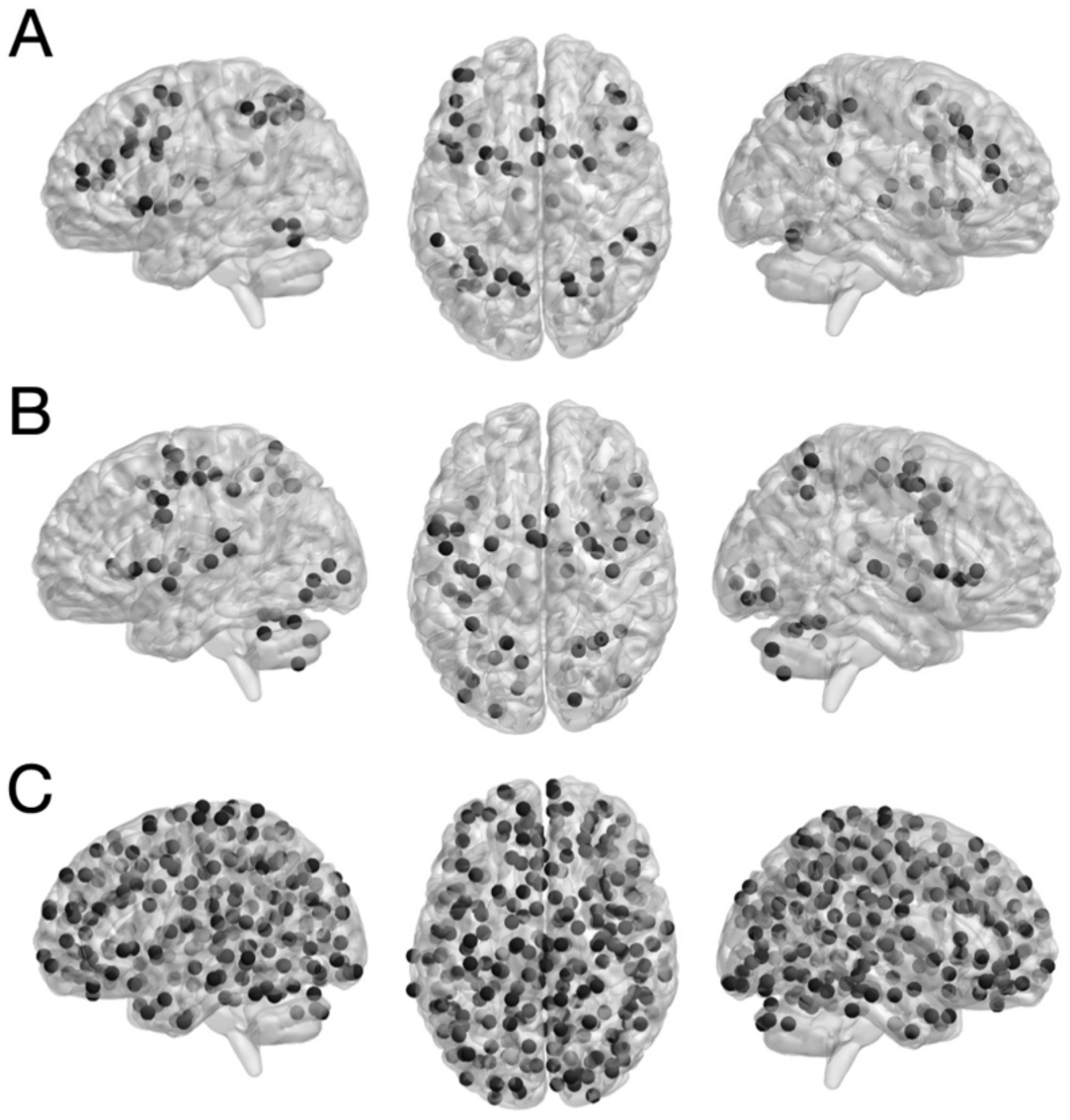
Nodes of meta-analytically defined A) executive function and B) perceptuo-motor networks, and C) Power et al.’s coordinates of putative functional areas. Taken with permission and modified from Heckner et al. (2023), copyright © 2023 Oxford University Press.

### Brain Metrics

Considering our multi-modal approach, we computed metrics for structural (GMV) and functional (RSFC, ReHo, and fALFF) modalities for every node in each network. Network nodes covered a sphere with 6-mm radius around each peak coordinate. Apart from the whole-brain network based on Power et al. (2011), peaks of meta-analytic convergence were extracted using the SPM Anatomy Toolbox version 3 (Eickhoff et al., 2005, 2007) and manually checked so that they would not overlap with each other or exceed the cortex when spheres were added. A gray-matter mask, including subcortical regions, was used to ascertain nodes comprised gray-matter (https://zenodo.org/record/6463123#.YlltJsjMJ3h).

### Gray-matter volume

Structural T1-weighted images were preprocessed and analyzed using SPM12 and the CAT12 toolbox. Within a unified segmentation model (Ashburner and Friston, 2000), images were corrected for bias-field inhomogeneities, brain tissue classified into gray matter, white matter, and cerebrospinal fluid, and spatially normalized to the Montreal Neurological Institute (MNI) template using DARTEL (Ashburner and Friston, 2011). Then, the segmented images were nonlinearly modulated using the Jacobian determinant derived from the normalization process to adjust them to the amount of expansion and contraction applied during normalization. GMV values were then obtained for each voxel in a given node as computed in CAT12 and then averaged across the node.

### Resting-state functional connectivity

The variance explained by the mean white-matter and cerebrospinal-fluid signal was removed from the time series to reduce spurious correlations. Subsequently, the data were band-pass filtered with the cut-off frequencies of 0.01 and 0.1 Hz.

There were no significant correlations between the target variables (i.e., task scores) and sex in either subgroup. The correlation between within-scanner movement (derivative of root mean square variance over voxels [DVARS]) and age was significant in the older subgroup. However, we refrained from additionally correcting for movement (i.e., removing movement-related variance that is partially shared with age). In our companion study, we additionally computed RSFC without global signal regression and with statistically removing the influence of the six head movement parameters (x, y, z translations and *α*, *β*, *γ* rotations) derived from realignment, their squared values as well as their derivatives to control for possible age-specific effects of global signal regression and movement. Importantly, both corrections did not alter the results, except for the main effect of age after movement correction, which was, however, qualified by the crossed age × task-demand level interaction and was therefore not interpreted. As such, movement regression may have removed age-related variance from the BOLD signal time series (Heckner et al., 2023) and was therefore not applied here.

In each network, RSFC was computed by first extracting the BOLD signal time-courses of all voxels within each network node expressed as the first eigenvariate. Then, pair-wise functional connectivity was computed as Fisher’s Z-transformed linear (Pearson) correlation between the first eigenvariate of the time series of each network’s nodes.

### Regional homogeneity

ReHo represents the homogeneity of a voxel’s time series with respect to its nearest local neighbors’ time courses (Zang et al., 2004) and is computed through Kendall’s coefficient of concordance (KCC; Kendall and Gibbons, 1990). Thus, each voxel is assigned a KCC value based on its time-series homogeneity towards its nearest neighbors and then averaged across the node.

### Fractional amplitude of low-frequency fluctuations

fALFF was computed as the ratio between power-spectrum in the frequency range (0.01 - 0.1 Hz) and spectral power in the entire frequency range. Therefore, the time series of each voxel was transformed to the frequency domain without band-pass filtering. Then, the square root was calculated at each frequency range of the power spectrum. Per voxel, the sum of power in the 0.01 - 0.1 Hz frequency range was divided by the summed power spectrum across the entire frequency range (Zou et al., 2008).

### Behavioral Measures

Executive function target variables were obtained from the eNKI-RS and comprised a high-demand (HD) and low-demand (LD; i.e., control) condition for each of three classical EF tasks (i.e., working memory, inhibitory control, and cognitive flexibility). All tasks used were previously evaluated and shown to have moderate to high reliability (Delis et al., 2001; Homack et al., 2005; Gur et al., 2010).

### Working memory

Working memory ability was quantified using reaction times (RT) of correct responses of the 1-back (HD) and 0-back (LD) conditions of the Short Letter-N-Back Test, which is part of Penn’s Computerized Neurocognitive Battery (CNB; Gur et al., 2010). In this test, participants are required to press a button if the letter on the screen is the same as the one presented N trials before.

### Inhibitory control

Inhibition performance was measured using RT of the incongruent (HD) and congruent (LD) conditions of the Color-Word Interference (CWI) Test, which is part of the Delis-Kaplan Executive Function System (D-KEFS; Delis et al., 2004). Here, participants are asked to name the ink color of a written word but inhibit the response to the word naming a color (same or different as the ink) itself.

### Cognitive flexibility

For quantifying cognitive flexibility, we used RT of the number and letter switching and sequencing conditions of the Trail Making Test (TMT), which is part of the D-KEFS. In this test, participants are asked to connect consecutive targets of one type (e.g., numbers; LD) or of two types (numbers and letters; HD) in an alternating fashion.

Raw performance scores for all tasks were *z*-transformed and outliers with more than three times the standard deviation below or above the mean were removed.

### Prediction

Individual *z*-transformed performance scores were then predicted from within-network GMV, RSFC, ReHo, and fALFF using partial least squares regression (PLSR; Krishnan et al., 2011). PLSR is similar to a supervised principal component regression (based on eigen-decomposition) and is thus advantageous when dimensionality reduction is beneficial for the analysis. In contrast to principal component regression, dimensionality reduction in PLSR is supervised (i.e., it uses information about the target variables), yielding the advantage that the resulting latent variables are all related to the target variables. After dimensionality reduction, a linear regressor was applied to the transformed data.

For prediction, a 10-fold cross-validation was performed for which the data were split into 10 sets, 9 of which were used for training while the 10^th^ was held back as a test set and subsequently used for prediction of the unseen data. This was done with each set being the test set once. In total, 100 repetitions of this 10-fold cross-validation were computed to ensure robustness. Prediction accuracy was assessed via RMSE, MAE, and Pearson’s *r*.

Prediction accuracy, as indicated by RMSE for the 100 repetitions, was then submitted to a 2 (age group) × 3 (network) × 2 (task-demand level) × 4 (modalities) mixed-measures ANOVA (*p* < .00005, Bonferroni-adjusted for the 10 × 100 cross-validation scheme), to further assess age differences in prediction accuracy, modality specificity, and the impact of task-demand level. Therefore, prediction results for low-demand (i.e., 0-back, CWI congruent, TMT consecutive) and high-demand (i.e., 1-back, CWI incongruent, TMT switch) conditions were averaged into LD and HD compound scores, respectively. When Mauchly’s test of sphericity was significant, Greenhouse-Geisser corrected results were interpreted.

To further corroborate the ANOVA main effects, machine-learning-adjusted *t*-tests for significant differences were computed (Nadeau and Bengio, 2003). To account for violating the independence assumption in a paired Student’s *t*-test, here, the variance estimate is adjusted by training and sample size.

## Results

### Prediction

The averaged prediction results from the test set as indicated by RMSE are displayed in Figure 2 and by Pearson’s *r* in Table 1. Additional accuracy measures, including RMSE and MAE, can be found in Tables S1-S6.

**Figure 2.**
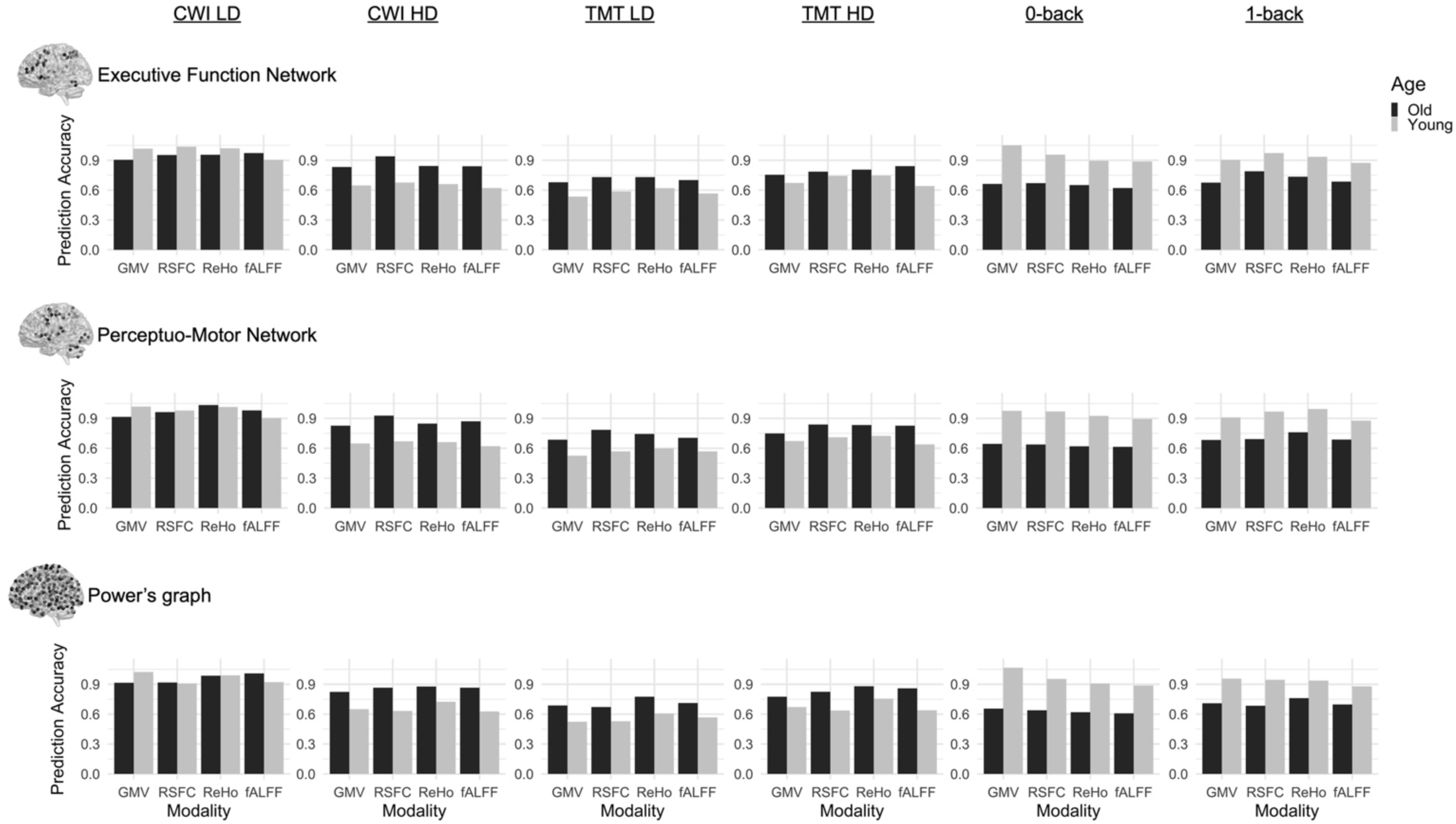
Prediction accuracies expressed as root mean squared error (RMSE) for Color Word Interference (CWI) low-demand (LD) congruent condition, CWI high-demand (HD) incongruent condition, Trail Making Test (TMT) LD consecutive condition, TMT HD switch condition, 0-back, and 1-back for old (dark) and young (light) adults from prediction within the executive function network, perceptuo-motor network, and Power’s graph of putative functional areas.

**Table 1.** Prediction Results as indicated by Pearson’s r according to Brain Modality, Brain Network and Target Variable for the Young and Old Subgroup.

|  |  | CWI_LD | CWI_HD | TMT_LD | TMT_HD | 0-back | 1-back |
| --- | --- | --- | --- | --- | --- | --- | --- |
| GMV |  |  |  |  |  |  |  |
| *EFN* | Old | .19 | .19 | .13 | .24 | .01 | .22 |
|  | Young | -.09 | .10 | .18 | -.01 | -.18 | .10 |
| *PercMot* | Old | .15 | .21 | .08 | .27 | .08 | .19 |
|  | Young | -.08 | .08 | .25 | -.02 | -.06 | .12 |
| *Power* | Old | .18 | .24 | .11 | .20 | .07 | .12 |
|  | Young | -.07 | .11 | .26 | .08 | -.16 | .02 |
| RSFC |  |  |  |  |  |  |  |
| *EFN* | Old | .10 | -.12 | .04 | .22 | .04 | -.13 |
|  | Young | -.04 | -.02 | .02 | -.27 | .10 | .01 |
| *PercMot* | Old | .01 | -.06 | -.21 | -.06 | .11 | .15 |
|  | Young | .05 | -.01 | .06 | -.22 | -.02 | -.04 |
| *Power* | Old | .13 | .03 | .11 | -.15 | -.05 | .10 |
|  | Young | .12 | .05 | .19 | .12 | -.15 | -.07 |
| ReHo |  |  |  |  |  |  |  |
| *EFN* | Old | .04 | .12 | -.07 | .07 | .05 | .02 |
|  | Young | -.04 | .12 | -.15 | -.26 | .12 | .12 |
| *PercMot* | Old | -.17 | .09 | -.07 | -.10 | .16 | -.18 |
|  | Young | -.07 | .10 | -.10 | -.22 | .01 | -.11 |
| *Power* | Old | .03 | .09 | -.07 | -.12 | .16 | -.04 |
|  | Young | -.01 | -.14 | -.02 | -.23 | .10 | .10 |
| fALFF |  |  |  |  |  |  |  |
| *EFN* | Old | -.02 | .06 | .01 | -.13 | .14 | .08 |
|  | Young | .06 | .09 | -.15 | 0 | .07 | .10 |
| *PercMot* | Old | -.06 | .04 | -.14 | -.04 | .18 | .05 |
|  | Young | .06 | .09 | -.18 | .03 | .04 | .06 |
| *Power* | Old | -.14 | .01 | .04 | -.15 | .18 | .05 |
|  | Young | .04 | .07 | -.10 | .04 | .07 | .08 |

*Note.* EFN = Executive Function Network, PercMot = Perceptuo-Motor Network, GMV = Gray-Matter Volume, RSFC = Resting-State Functional Connectivity, ReHo = Regional Homogeneity, fALFF = Fractional Amplitude of Low-Frequency Fluctuations, CWI = Color-Word Interference, TMT = Trail Making Test, LD = Low-Demand, HD = High-Demand.

### Mixed-Measures ANOVA

To further assess age differences in prediction accuracy as well as network specificity and the impact of the task-demand level, we submitted the prediction accuracies as given by RMSE values for each of the 100 repetitions to a 2 (age group) × 3 (network) × 2 (task-demand level) × 4 (modalities) mixed-measures ANOVA (*p* < .00005, Bonferroni-adjusted for the 10 × 100 cross-validation scheme).

In a first step, prediction results for low-demand (i.e., 0-back, CWI congruent, TMT consecutive) and high-demand (i.e., 1-back, CWI incongruent, TMT switch) conditions were averaged into LD and HD compound scores, respectively. The ANOVA yielded significant main effects for the factors modality, network, task-demand level, and age group. These effects were qualified by two-way, three-way, and a four-way interaction among all factors (see Table 2).

**Table 2.**
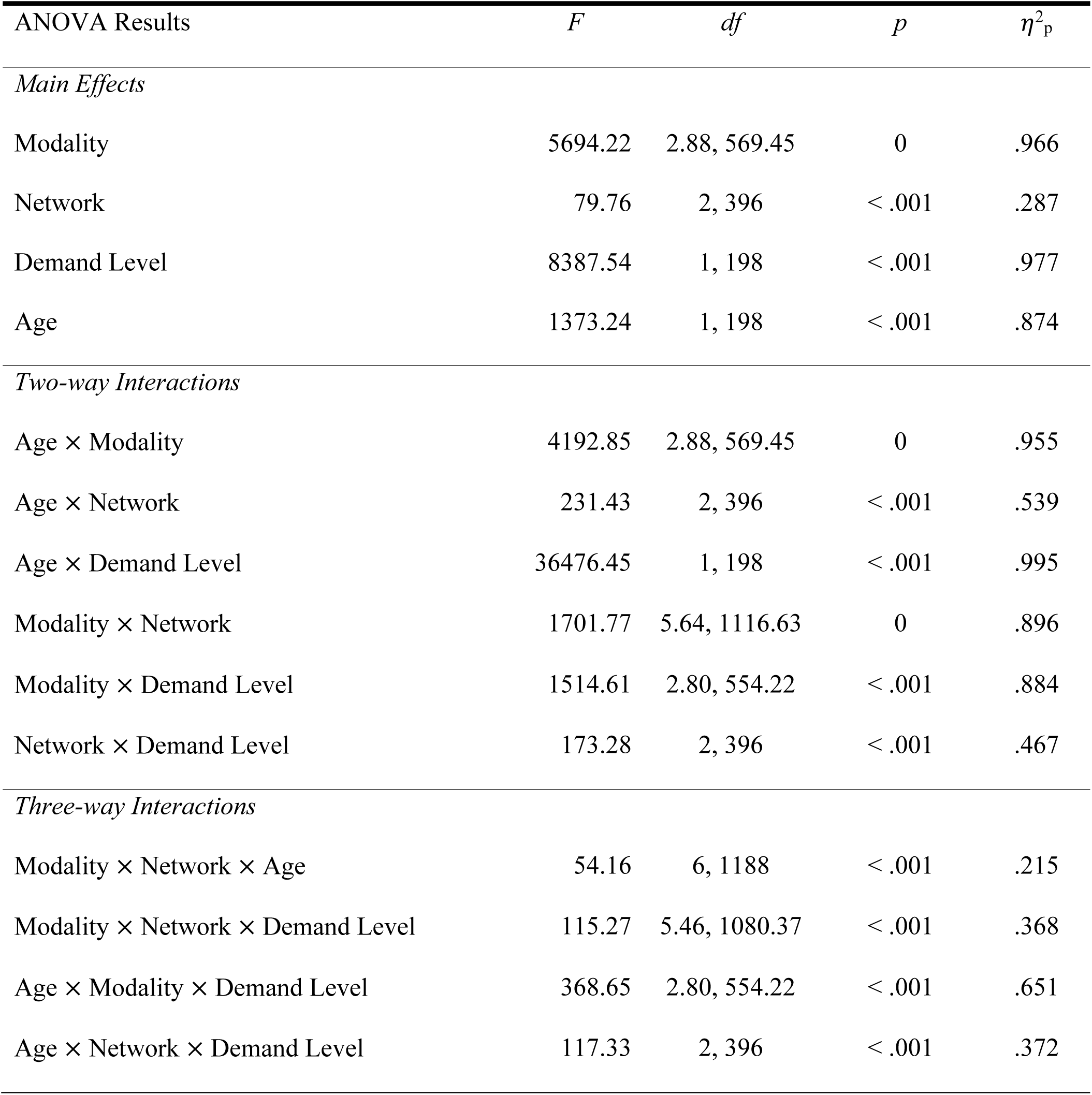
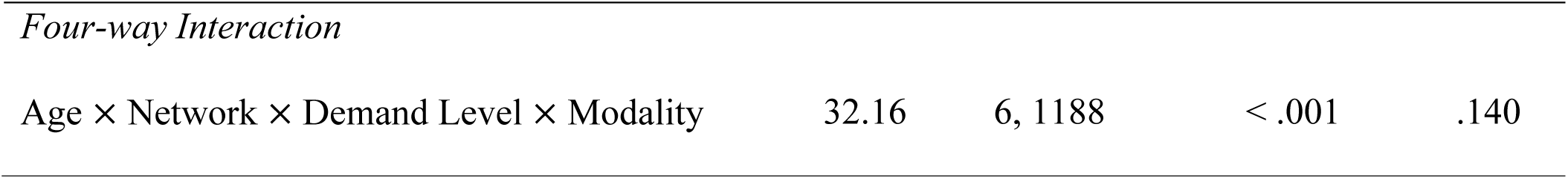
ANOVA Results.

| ANOVA Results | *F* | *df* | *p* | $\eta^2_p$ |
| --- | --- | --- | --- | --- |
| *Main Effects* |  |  |  |  |
| Modality | 5694.22 | 2.88, 569.45 | 0 | .966 |
| Network | 79.76 | 2, 396 | < .001 | .287 |
| Demand Level | 8387.54 | 1, 198 | < .001 | .977 |
| Age | 1373.24 | 1, 198 | < .001 | .874 |
| *Two-way Interactions* |  |  |  |  |
| Age × Modality | 4192.85 | 2.88, 569.45 | 0 | .955 |
| Age × Network | 231.43 | 2, 396 | < .001 | .539 |
| Age × Demand Level | 36476.45 | 1, 198 | < .001 | .995 |
| Modality × Network | 1701.77 | 5.64, 1116.63 | 0 | .896 |
| Modality × Demand Level | 1514.61 | 2.80, 554.22 | < .001 | .884 |
| Network × Demand Level | 173.28 | 2, 396 | < .001 | .467 |
| *Three-way Interactions* |  |  |  |  |
| Modality × Network × Age | 54.16 | 6, 1188 | < .001 | .215 |
| Modality × Network × Demand Level | 115.27 | 5.46, 1080.37 | < .001 | .368 |
| Age × Modality × Demand Level | 368.65 | 2.80, 554.22 | < .001 | .651 |
| Age × Network × Demand Level | 117.33 | 2, 396 | < .001 | .372 |
| *Four-way Interaction* |  |  |  |  |
| Age × Network × Demand Level × Modality | 32.16 | 6, 1188 | < .001 | .140 |

The complementary machine-learning-adjusted *t*-tests conducted to corroborate the main effects did not confirm the main effects of age and task-demand level. Network differences were only significant for perceptuo-motor vs. Power networks, but not for EF vs. perceptuo-motor and EF vs. Power networks, respectively. All modality differences but fALFF vs. GMV were significant (see Table S7).

We obtained the following results from the two-way interactions. While prediction accuracy in older participants was significantly better for LD than HD task conditions, prediction accuracy in younger participants was better for HD than LD conditions (see Figure 3A and Table 3). Regarding network differences, prediction accuracy for LD conditions was best for the whole-brain approach, as compared to the perceptuo-motor network and the EFN, whereas for HD conditions, it was best for the EFN (see Figure 3B and Table 3). Furthermore, in older adults, prediction accuracy was best for the EFN, relative to the whole-brain and perceptuo-motor networks. Conversely, for younger adults, prediction accuracy was generally best for the whole-brain network (see Figure 3C and Table 3).

**Figure 3.**
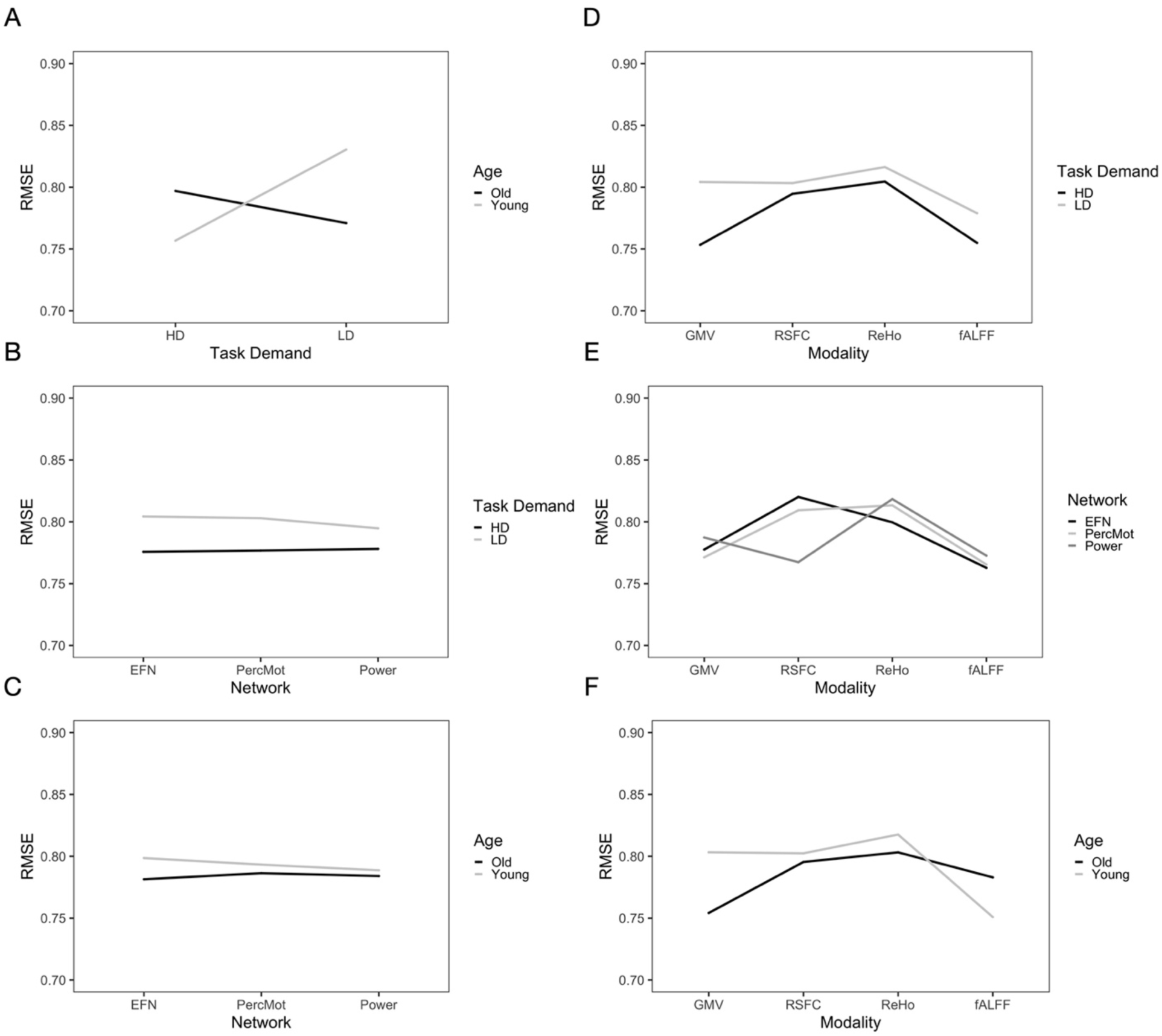
Interaction effects for (A) age × demand level, (B) demand level × network, (C) age × network, (D) demand level × modality, (E) network × modality, and (F) age × modality.

**Table 3.**
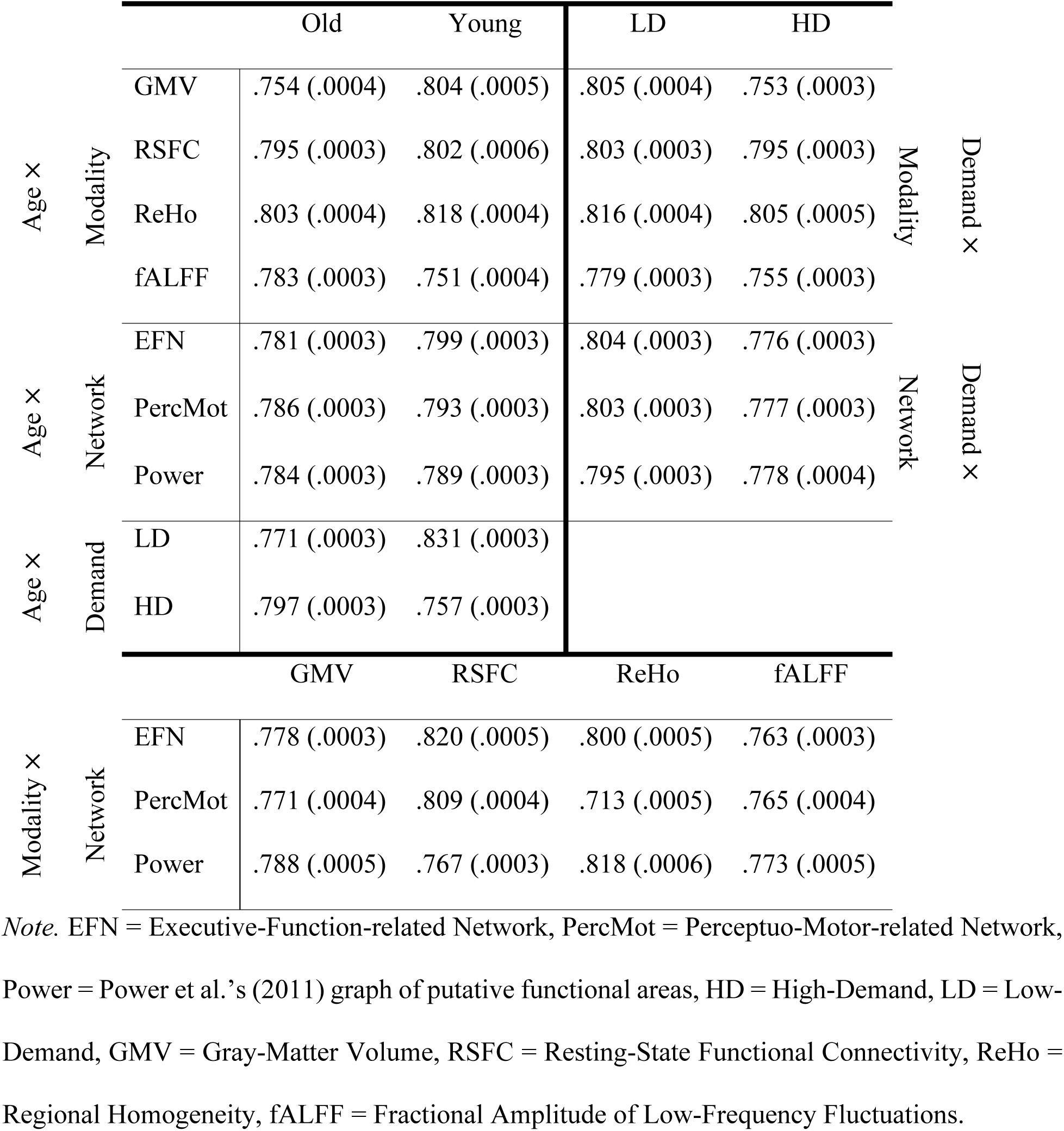
ANOVA Interaction Effects Displayed as Mean and Standard Error.

For LD conditions, prediction accuracy was generally best for fALFF, as compared to RSFC, GMV, and ReHo, while for HD conditions, prediction accuracy was generally best for GMV (see Figure 3D and Table 3). For ReHo and fALFF, prediction accuracy was generally best for the EFN, while for RSFC, prediction accuracy was best for the whole-brain network, and for GMV for the perceptuo-motor network (see Figure 3E and Table 3). For older adults, prediction accuracy was generally the highest for GMV, as compared to fALFF, RSFC, and ReHo, while for younger adults, prediction accuracy was best for fALFF (see Figure 3F and Table 3).

Post-hoc pairwise comparisons revealed that prediction accuracy (i.e., RMSE) was better for older than younger participants (see Figure 4A and Table 4). Prediction accuracy was better for HD as compared to LD (see Figure 4B and Table 4) conditions across networks, modalities, and age groups. Across demand level, networks, and age groups, prediction accuracy was best for fALFF as compared to GMV, RSFC, and ReHo (see Figure 4C and Table 4). Prediction accuracy was best for the Power nodes as compared to the EFN, and perceptuo-motor network (see Figure 4D and Table 4). The EFN and perceptuo-motor network did not differ significantly.

**Figure 4.**
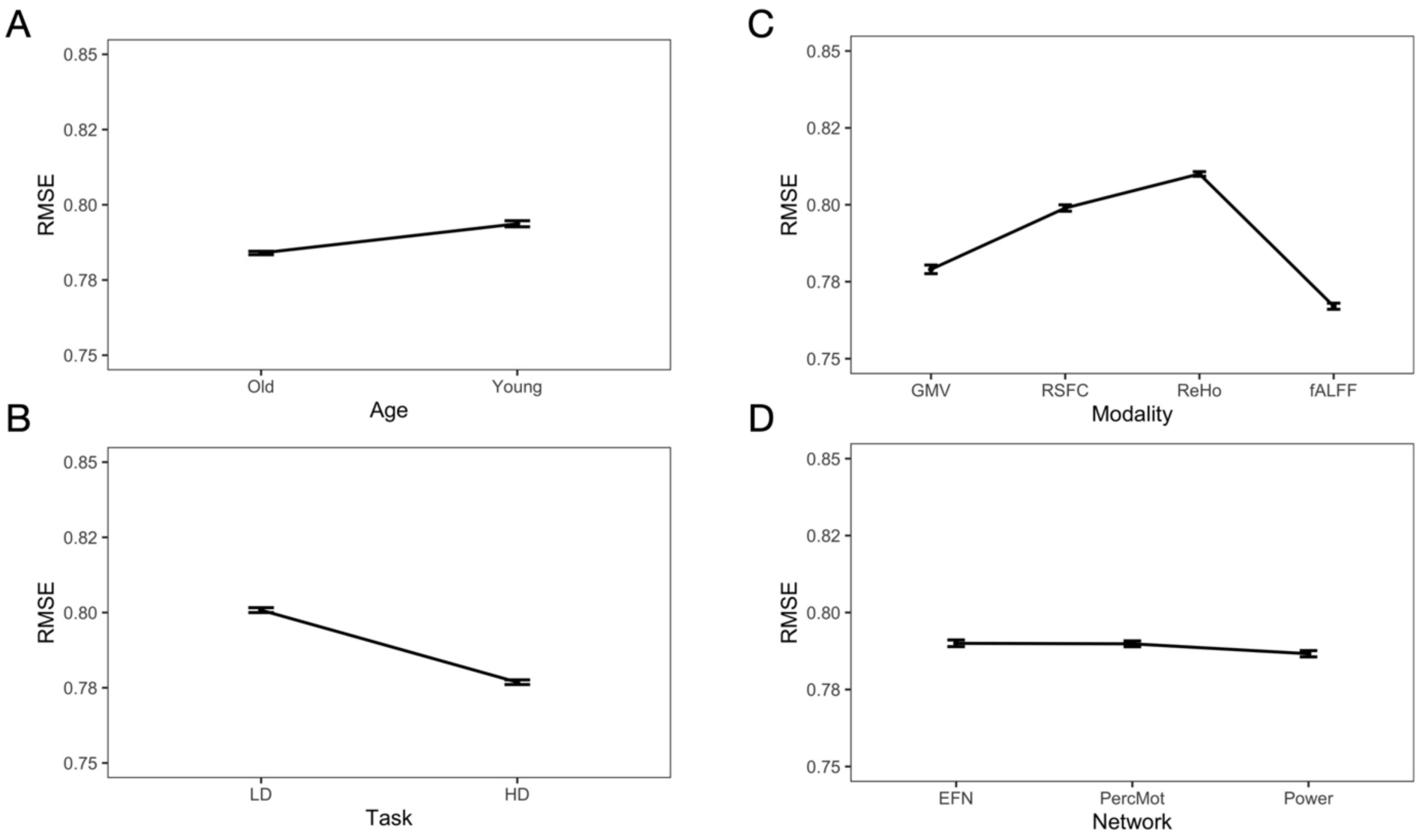
Main effect for age, task, modality, and network (mean ± standard error).

**Table 4.** Post-hoc Pairwise Comparisons of ANOVA Effects.

| Post-hoc Pairwise Comparisons |  |  |  |
| --- | --- | --- | --- |
| Factor | Mean (SE)<sub>group1</sub> | Mean (SE)<sub>group2</sub> | *p* |
| Age | .784 (.0002)<sub>old</sub> | .794 (.0002)<sub>young</sub> | $5.30 \times 10^{-91}$ |
| Modality | .779 (.0002)<sub>GMV</sub> | .767 (.0002)<sub>fALFF</sub> | $3.27 \times 10^{-87}$ |
| | .779 (.0002)<sub>GMV</sub> | .799 (.0002)<sub>RSFC</sub> | $4.33 \times 10^{-129}$ |
| | .779 (.0002)<sub>GMV</sub> | .810 (.0003)<sub>ReHo</sub> | $7.67 \times 10^{-153}$ |
| | .799 (.0002)<sub>RSFC</sub> | .767 (.0002)<sub>fALFF</sub> | $3.75 \times 10^{-166}$ |
| | .799 (.0002)<sub>RSFC</sub> | .810 (.0003)<sub>ReHo</sub> | $3.27 \times 10^{-74}$ |
| | .767 (.0002)<sub>fALFF</sub> | .819 (.0003)<sub>ReHo</sub> | $4.48 \times 10^{-161}$ |
| Network | .790 (.0002)<sub>EFN</sub> | .790 (.0002)<sub>PercMot</sub> | .583 |
| | .790 (.0002)<sub>EFN</sub> | .787 (.0002)<sub>Power</sub> | $5.01 \times 10^{-25}$ |
| | .790 (.0002)<sub>PercMot</sub> | .787 (.0002)<sub>Power</sub> | $1.24 \times 10^{-25}$ |
| Demand Level | .777 (.0002)<sub>HD</sub> | .801 (.0002)<sub>LD</sub> | $4.84 \times 10^{-180}$ |

*Note.* GMV = Gray-Matter Volume, RSFC = Resting-State Functional Connectivity, ReHo = Regional Homogeneity, fALFF = Fractional Amplitude of Low-Frequency Fluctuations, EFN = Executive-Function-related Network, PercMot = Perceptuo-Motor-related Network, Power = Power et al.'s (2011) graph of putative functional areas, HD = High-Demand, LD = Low-Demand, SE = Standard Error.

## Discussion

The current study investigated to what extent regional brain morphology (GMV) and three different functional brain metrics (RSFC, ReHo, and fALFF) predict individual differences in EF abilities, and if these brain–behavior associations are modality- and/or age-specific. For this purpose, we defined three brain networks: an EF-related network, a perceptuo-motor network linked to visual, auditory, and motor processing, and a whole-brain network. We predicted individual EF performance scores of three critical EF subcomponents (i.e., working memory, inhibitory control, and cognitive flexibility) from within-network GMV, RSFC, ReHo, and fALFF. Finally, we submitted the prediction results to a 2 (age group) × 4 (modalities) × 3 (network) × 2 (task-demand level) mixed-measures ANOVA to assess the effects of modality and age. While prediction accuracy was overall rather low to moderate, it was better for high-than low-demand task conditions. This difference was especially pronounced for fALFF and GMV. However, this effect might be driven by the age × task-demand level interaction, as prediction accuracy for younger adults was better for HD (vs. LD), whereas for older adults, it was better for LD (vs. HD) conditions. Prediction accuracy for younger adults was best with fALFF, while for older adults, the highest accuracy was achieved with GMV.

### Prediction of EF Abilities

In line with our companion study (Heckner et al., 2023) as well as published guidelines for prediction analyses (Scheinost et al., 2019; Poldrack et al., 2020), we assessed our prediction results with RMSE, MAE, as well as Pearson’s correlation coefficient (*r*) as these scores offer different, yet complementary, information about the accuracy of predictive models and the association between brain metrics and behavioral target variables. Here, we will discuss prediction accuracy as measured with RMSE (< .8) and the respective Pearson’s *r* correlation coefficient (note that cognitive performance was *z*-scored). As mentioned above, prediction accuracies and brain–behavior associations were moderate at best, and this was also evident from the explained variance of the prediction models, as measured with the coefficient of determination R^2^ (Scheinost et al., 2019). In our study, the explained variance did not exceed 6%. Whole-brain RSFC was, for example, only able to explain 3.7% of the variance in the TMT LD condition for younger adults (*r* = .19). GMV within the EFN was able to explain 3.7% of the variance in the TMT HD condition for older adults (*r* = .24). Similarly, but using the perceptuo-motor network, 5.2% variance was explained for the same target for older adults (*r* = .27) and only 2.1% targeting TMT LD condition in the young (*r* = .25). Additionally, using GMV from whole-brain features, 1.5% variance in TMT HD condition was explained for older adults (*r* = .20) and 2.7% in the TMT LD condition for the young (*r* = .26).

Predictions based on ReHo were not able to explain any variance in the target variables. Within-EFN fALFF, on the other hand, explained 2.1% of variance in the working memory LD condition for older adults (*r* = .14) and 2.6% when predicting from the whole-brain (*r* = .18). From these results, it is surprising to note that, while prediction from within-network fALFF resulted in the best overall prediction accuracy, only very little variance could be eventually explained. GMV was able to explain variance in more tasks and conditions. Nevertheless, the amount of explained variance was overall still quite low (R^2^ < 0.06). Thus, the question arises how to harmonize the finding of better prediction accuracies but quite low brain–behavior associations for features extracted from fALFF. One possibility would be that within-subject functional brain variability may be highly important, even necessary, for EF but that interindividual differences in this variability, as reflected by fALFF, do not scale with individual EF abilities, at least not in the normal range of performance. More work is needed to understand the neural mechanisms and functional meaning of fALFF.

Summing up, age group, as well as task and demand levels, show moderate modality specificity. Brain–behavior associations were generally rather low but more pronounced when predicting from structural as compared to functional features. These findings bring into question whether it is feasible to predict individual EF abilities from functional metrics at rest as well as from a priori brain networks defined via group-level analyses. Together with recent evidence from RSFC-based predictive modelling (Heckner et al., 2023), the present results argue against network specificity, but this time even across different brain feature modalities. Importantly, the results across both of our studies stress the need for brain measures that are not just somewhat associated with EF but can actually explain variance in individual EF abilities. While brain– behavior associations found in the present study are generally low, they are comparable to other results in the field (Ferguson et al., 2017; Greene et al., 2018; He et al., 2021), calling for more informative measures or methods and for a critical re-evaluation of the predictive and explanatory value of the models examined so far.

### Modality Specificity and Age Effects

The ANOVA yielded a main effect of modality on prediction accuracy, and post-hoc pairwise comparisons revealed significant differences between all modalities. Overall, the best prediction accuracy was achieved when predicting from fALFF, followed, in descending order, by GMV, RSFC, and ReHo. However, this main effect may be explained by the age × modality interaction. While for younger adults, prediction was best from fALFF, followed, in descending order, by GMV, RSFC, and ReHo, for older it was best from GMV, followed by RSFC, fALFF, and ReHo. RSFC’s better prediction accuracy, however, may be explained by the modality × network interaction. This interaction revealed that prediction from RSFC was best for the whole-brain approach. One possible explanation for this finding might be the greater feature space gain for the whole-brain approach with 34,716 connections as compared to 1,225 connections for the EFN. However, in our companion study (Heckner et al., 2023), 10 random networks of the same size as the EFN still resulted in a significantly better prediction accuracy than the EFN and the perceptuo-motor network. Therefore, it does not seem to be the sheer number of features that is responsible for the prediction outcome. Lastly, all metrics were better at predicting HD (vs. LD) task conditions. This effect was especially pronounced for features extracted from GMV and fALFF. Again, this main effect is qualified by the age × demand level interaction, which revealed best prediction accuracies for older adults for LD conditions, whereas for younger adults, best prediction accuracies were achieved for HD conditions. As such, the main effect of task demand appears to be driven by older adults and should not be interpreted because of the underlying crossed interaction.

Our results revealed that GMV and fALFF contained more information on individual EF performance than did the other modalities and that this effect was age-dependent. Regional GMV is very well linked to global atrophy observed in advanced age. Previous research, however, has shown that the age-related decline in GMV is especially pronounced in brain regions associated with EF, such as fronto-parietal areas (Taki et al., 2004; Chee et al., 2006; Hu et al., 2014). Furthermore, the pattern of global decline is thought to be rather consistent across older adults (bilateral pre-supplementary motor area [pre-SMA], supplementary motor area [SMA], insula, anterior cingulate cortex [ACC], dorsolateral prefrontal cortex [DLPFC], inferior parietal lobule [IPL], and caudate; Bergfield et al., 2010; Giorgio et al., 2010; Taki et al., 2011). In line with these findings, the current results suggest that GMV may be a possible marker for individual EF ability levels in older adults – although one must keep in mind that prediction accuracies were still only small to moderate.

For younger adults, the best prediction accuracy was achieved with fALFF. Previous research suggested that low-frequency fluctuations of the BOLD signal are spontaneous and reflect the intrinsic connectivity of the brain (Biswal et al., 1995; Fox and Raichle, 2007). Thus, fALFF is understood as a measure of functional within-subject variability that possibly reflects cognitive adaptability (i.e., the ease of mental set reconfiguration) to task demands (Bolt et al., 2018; Uddin, 2020). In young adults, the fALFF pattern appears to be linked to behavior more closely, while in older adults, fALFF does not seem to contain relevant information about individual EF performance. Previous research has shown that an age-related decrease in fALFF and GMV overlapped in prefrontal/frontal brain regions including pre-SMA, SMA, and DLPFC (Hu et al., 2014). It was concluded that prefrontal brain regions, critical for EF, show concurrent age-related changes in structure and function. Earlier it had been suggested that younger, faster participants show a higher variability in brain activity across tasks and greater regional dedifferentiation of signal variability than older adults (Garrett et al., 2011). As variability may provide the kinetic energy for brain networks to explore possible functional architectures (McIntosh et al., 2010; Deco et al., 2011), an intrinsically more variable brain might be able to configure optimal networks for processing a given input towards a particular behavioral goal more flexibly and efficiently (Garrett et al., 2011).

One reason for the rather low brain–behavior associations achieved from within-network RSFC might be its unconstrained nature, as discussed in detail in our companion paper (Heckner et al., 2023). Several recent studies have shown that behavioral prediction from brain connectivity during tasks (or movie watching) may work somewhat better than from rest (Greene et al., 2018; Sripada et al., 2020; Finn and Bandettini, 2021; Kraljević et al., 2023). Tasks modulate functional brain states and may thus offer important information about individual differences in brain functional organization and their association with behavior (Greene et al., 2018). ReHo, a measure of local synchronicity/connectivity, is thought to induce global connectivity (i.e., RSFC; Jiang and Zuo, 2016). Therefore, it would not be surprising if both local and global connectivity measures may be affected by, for example, mind wandering or thinking about a task during rest (Gregory et al., 2016). fALFF, on the other hand, as a measure of local variability or adaptability, reflecting the spontaneous, intrinsic connectivity of the brain, may therefore be influenced to a lesser degree by unconstrained thoughts during rest. Such a differential susceptibility to state effects might possibly explain the lower prediction accuracies observed for RSFC and ReHo and the superiority of fALFF, even though all metrics are based on brain activity during “rest” (i.e., in a state without an externally driven task).

Differential susceptibility to state effects might have also contributed to the relatively better prediction performance observed for within-network GMV, which was best for older adults and second best for younger adults, as compared to the other brain metrics. In particular, functional metrics might be overall more susceptible to state effects and, thus, have a reliability disadvantage, relative to structural metrics. Hence, lower prediction accuracies achieved for the functional metrics, especially RSFC and ReHo, should not prematurely be marked as weak markers for any individual performance differences but rather as a product of state–trait interactions – which would make these metrics less suitable for capturing stable, trait-level aspects of brain activity that are thought to share variance with stable cognitive traits, in particular when the amount of brain activity data available per participant is relatively limited. For more conclusive answers, future studies are needed that investigate the comparative reliability of different functional and structural brain metrics a well as the impact of age on this issue (see, e.g., Song et al., 2012, for reporting on age-related RSFC reliability differences).

Similar to previous studies (Pläschke et al., 2020; Heckner et al., 2023), prediction accuracy across modalities, networks, and task-demand levels was better for older (vs. younger) subjects, suggesting that brain–behavior associations become tighter with advancing age. Possibly, such increased associations might be due to overall age-related neural decline, such as brain atrophy or white-matter degeneration, influencing network integrity (Cabeza et al., 2016) and reorganization that is linked to EF performance. However, this main effect is qualified by the crossed age × demand level interaction and should thus only be interpreted with great caution.

Interestingly, we replicated this age × demand level interaction already reported in Heckner et al. (2023) across all structural and functional modalities, such that prediction accuracy for younger adults was consistently better when predicting HD (vs. LD) conditions, while for older adults, the reverse pattern was observed. One possible explanation is that age-related effects on the network-level might still be compensated for in low-demand conditions, but they might not in high-demand conditions taxing EF abilities. This is in line with the compensation-related utilization of neural circuits hypothesis of cognitive aging (Reuter-Lorenz and Cappell, 2008) and previous research showing that age-related neurobiological decline comprises BOLD responsivity during task state. Older adults might be able to compensate LD task conditions but reach a ceiling at a certain level of processing demands such that compensatory activation cannot be further increased in HD conditions (Nagel et al., 2011). Hence, our results emphasize the relevance of behavioral testing procedures that are more adaptive to performance differences (e.g., because of compensatory efforts) in order to have tests that are sensitive enough to capture meaningful brain–behavior associations across ability levels (cf. Heckner et al., 2023). Interestingly, our results revealed a trend toward better prediction accuracies as well as stronger brain–behavior associations for TMT scores, as compared to CWI and n-back performance, across demand level, networks, and age groups for features extracted from GMV and RSFC. Although we did not specifically aim to investigate prediction performance depending on the individual EF tasks, the TMT might be a more sensitive target variable in capturing individual differences in EF performance as compared to the other tests used. Possibly, because the TMT measures different facets and stages of cognitive processing and might therefore be sensitive to several changes in cognition. This might especially be the case, when targeting LD and HD conditions separately and not subtracted (i.e., HD-LD).

Overall, despite generally low prediction accuracies, our results point out the superiority of GMV and fALFF in predicting individual differences in EF performance, and indicate that this effect is age dependent. While older adults’ EF performance was predicted best by features extracted from GMV, younger adults’ EF performance was predicted best by fALFF features. For older adults, overall patterns, like global atrophy, seem to be most predictive of EF performance in comparison to the other metrics applied. For younger adults, fALFF appears to be most predictive of EF performance. As a measure of neural variability, fALFF may reflect the ease of exploring the best suited network constellation for a given task. Additionally, our results stress the importance of adaptive testing in order to find meaningful brain–behavior associations.

### Conclusion and Outlook

The current study investigated to what extent individual differences in EF performance can be predicted from structural (GMV), as well as resting-state functional (RSFC, ReHo, and fALFF) modalities. In addition, we examined whether the pattern in prediction changes with age, brain network, or task-demand level. Our results revealed overall rather low to moderate prediction accuracies and brain–behavior associations. Explained variance in the target variables did not exceed 6%. These findings generally question the utility of the brain metrics examined here for predicting individual differences in EF abilities. However, our results did point out the superiority of GMV and fALFF as compared to ReHo and RSFC in predicting individual EF performance. One possibility could be that individual differences in EF abilities are more strongly driven by global brain characteristics that can be better assessed with metrics for global atrophy or variability. Furthermore, our results revealed an age-related modality specificity: For older adults, structural measures of overall atrophy might be more informative, while for younger adults, functional measures of brain variability seem to contain more information about individual EF abilities.

The overall low to moderate prediction accuracy as well as the missing network specificity questions the potential of the single metrics to be applied as biomarkers for individual differences in EF performance. Rather, our findings suggest that future research may need to analyze more global properties of the brain, possibly combining different structural and functional metrics, to result in more sensitive predictors for young and older adults, respectively. This also applies to adaptive behavioral testing as our results revealed better prediction accuracies in LD (vs. HD) task conditions for older adults, while for younger adults, prediction accuracies were better for HD (vs. LD) task conditions. Furthermore, it is important to consider the possible impact of the feature space size, especially when comparing different metrics (e.g., edge-level RSFC vs. node-level ReHo). A replication with a different, larger sample as well as different cognitive states (i.e., task performance, movie watching) and a continuous age distribution might prove useful for revealing more information contained in the brain about individual mental abilities. Lastly, considering the complexity of machine learning outputs and the increased use and relevance of these approaches in the field of behavioral neuroscience, developing appropriate methods for comparing the outcomes of different models accounting for intrinsic dependencies in the cross-validation scheme is strongly warranted.

## Competing interests

The authors declare that the research was conducted in the absence of any commercial or financial relationships that could be construed as a potential conflict of interest.

## Acknowledgements

This study was supported by the Deutsche Forschungsgemeinschaft (DFG, EI 816/11-1, PA 3634/1-1, EI 816/21-1, SPP2041, CRC), the National Institute of Mental Health (R01-MH074457), the Helmholtz Portfolio Theme “Supercomputing and Modeling for the Human Brain”, the European Union’s Horizon 2020 Research and Innovation Programme under Grant Agreement No. 720270 (HBP SGA1), 785907 (HBP SGA2), and 945539 (HBP SGA3).

## Notes

### Competing Interest Statement

The authors have declared no competing interest.

