## Supplementary material for "Predicting Executive Functioning from Brain Networks: Modality Specificity and Age Effects": Tables S1-S6; Table S7

Table S1

*Prediction Accuracies for Low-demand Color-Word Interference Task Performance in the Young and Old Subgroup*

| GMV | | $r_{old}$ | $r_{young}$ | $MAE_{old}$ | $MAE_{young}$ | $RMSE_{old}$ | $RMSE_{young}$ |
|---|---|---|---|---|---|---|---|
| | EFN | .19 | -.09 | .68 | .81 | .9 | 1.02 |
| | PercMot | .15 | -.08 | .7 | .81 | .92 | 1.02 |
| | Power | .18 | -.07 | .7 | .81 | .92 | 1.02 |
| RSFC | | | | | | | |
| | EFN | .1 | -.04 | .74 | .85 | .95 | 1.04 |
| | PercMot | .01 | .05 | .75 | .76 | .96 | .98 |
| | Power | .13 | .12 | .72 | .74 | .92 | .91 |
| ReHo | | | | | | | |
| | EFN | .04 | -.04 | .73 | .82 | .91 | 1.02 |
| | PercMot | -.17 | -.07 | .82 | .81 | 1.03 | 1.01 |
| | Power | .03 | -.01 | .76 | .8 | .99 | .99 |
| fALFF | | | | | | | |
| | EFN | -.02 | .06 | .74 | .72 | .97 | .9 |
| | PercMot | -.06 | .06 | .76 | .72 | .98 | .9 |
| | Power | -.14 | .04 | .78 | .73 | 1.01 | .92 |

*Note.* MAE = mean absolute error, RMSE = root mean squared error, EFN = executive function network, PercMot = perceptuo-motor network, GMV = gray-matter volume, RSFC = resting-state functional connectivity, ReHo = regional homogeneity, fALFF = fractional amplitude of low frequency fluctuations.

Table S2

*Prediction Accuracies for High-demand Color-Word Interference Task Performance in the Young and Old Subgroup*

| GMV | | $r_{old}$ | $r_{young}$ | $MAE_{old}$ | $MAE_{young}$ | $RMSE_{old}$ | $RMSE_{young}$ |
|---|---|---|---|---|---|---|---|
| | EFN | .19 | .1 | .67 | .52 | .83 | .65 |
| | PercMot | .21 | .08 | .66 | .53 | .83 | .65 |
| | Power | .24 | .11 | .66 | .52 | .82 | .65 |
| RSFC | | | | | | | |
| | EFN | -.12 | -.02 | .74 | .56 | .94 | .68 |
| | PercMot | -.06 | -.01 | .72 | .55 | .93 | .67 |
| | Power | .03 | .05 | .67 | .53 | .87 | .63 |
| ReHo | | | | | | | |
| | EFN | .12 | .12 | .66 | .53 | .84 | .66 |
| | PercMot | .09 | .1 | .66 | .54 | .85 | .66 |
| | Power | .09 | -.14 | .69 | .59 | .88 | .72 |
| fALFF | | | | | | | |
| | EFN | .06 | .09 | .65 | .51 | .84 | .62 |
| | PercMot | .04 | .09 | .68 | .51 | .87 | .62 |
| | Power | .01 | .07 | .67 | .52 | .87 | .63 |

*Note.* See Table S1.

Table S3

*Prediction Accuracies for Low-demand Trail Making Test Performance in the Young and Old Subgroup*

| GMV | | $r_{old}$ | $r_{young}$ | $MAE_{old}$ | $MAE_{young}$ | $RMSE_{old}$ | $RMSE_{young}$ |
|---|---|---|---|---|---|---|---|
| | EFN | .13 | .18 | .54 | .41 | .46 | .53 |
| | PercMot | .08 | .25 | .53 | .41 | .69 | .53 |
| | Power | .11 | .26 | .54 | .41 | .69 | .52 |
| RSFC | | | | | | | |
| | EFN | .04 | .02 | .57 | .47 | .73 | .59 |
| | PercMot | -.21 | .06 | .63 | .46 | .79 | .57 |
| | Power | .11 | .19 | .53 | .42 | .67 | .53 |
| ReHo | | | | | | | |
| | EFN | -.07 | -.15 | .57 | .48 | .73 | .62 |
| | PercMot | -.07 | -.1 | .59 | .47 | .74 | .6 |
| | Power | -.07 | -.02 | .62 | .48 | .78 | .61 |
| fALFF | | | | | | | |
| | EFN | .01 | -.15 | .56 | .44 | .7 | .57 |
| | PercMot | -.14 | -.18 | .56 | .44 | .71 | .57 |
| | Power | .04 | -.1 | .58 | .44 | .71 | .57 |

*Note.* See Table S1.

Table S4

*Prediction Accuracies for High-demand Trail Making Test Performance in the Young and Old Subgroup*

| GMV | | $r_{old}$ | $r_{young}$ | $MAE_{old}$ | $MAE_{young}$ | $RMSE_{old}$ | $RMSE_{young}$ |
|---|---|---|---|---|---|---|---|
| | EFN | .24 | -.01 | .57 | .51 | .75 | .67 |
| | PercMot | .27 | -.02 | .57 | .52 | .75 | .67 |
| | Power | .2 | .08 | .58 | .51 | .77 | .67 |
| RSFC | | | | | | | |
| | EFN | .22 | -.27 | .62 | .59 | .78 | .74 |
| | PercMot | -.06 | -.22 | .65 | .54 | .84 | .71 |
| | Power | -.15 | .12 | .62 | .50 | .82 | .64 |
| ReHo | | | | | | | |
| | EFN | .07 | -.26 | .61 | .58 | .81 | .75 |
| | PercMot | -.1 | -.22 | .64 | .57 | .83 | .72 |
| | Power | -.12 | -.23 | .67 | .58 | .88 | .76 |
| fALFF | | | | | | | |
| | EFN | -.13 | 0 | .63 | .51 | .84 | .64 |
| | PercMot | -.04 | .03 | .64 | .5 | .83 | .64 |
| | Power | -.15 | .04 | .65 | .5 | .86 | .64 |

*Note.* See Table S1.

Table S5

*Prediction Accuracies for Low-demand N-Back Test Performance in the Young and Old Subgroup*

| GMV | | $r_{old}$ | $r_{young}$ | $MAE_{old}$ | $MAE_{young}$ | $RMSE_{old}$ | $RMSE_{young}$ |
|---|---|---|---|---|---|---|---|
| | EFN | .01 | -.18 | .53 | .82 | .66 | 1.1 |
| | PercMot | .08 | -.06 | .51 | .75 | .64 | .98 |

| | | | | | | | |
|---|---|---|---|---|---|---|---|
| | Power | .07 | -.16 | .52 | .82 | .66 | 1.07 |
| RSFC | | | | | | | |
| | EFN | .04 | .10 | .54 | .74 | .67 | .96 |
| | PercMot | .11 | -.02 | .51 | .76 | .64 | .97 |
| | Power | -.05 | -.15 | .51 | .74 | .64 | .95 |
| ReHo | | | | | | | |
| | EFN | .05 | .12 | .52 | .7 | .65 | .89 |
| | PercMot | .16 | .01 | .49 | .72 | .62 | .93 |
| | Power | .16 | .1 | .5 | .72 | .62 | .91 |
| fALFF | | | | | | | |
| | EFN | .14 | .07 | .49 | .69 | .62 | .89 |
| | PercMot | .18 | .04 | .48 | .69 | .61 | .89 |
| | Power | .18 | .07 | .48 | .69 | .61 | .89 |

*Note.* See Table 1.

Table S6

*Prediction Accuracies for High-demand N-Back Test Performance in the Young and Old Subgroup*

| GMV | | $r_{old}$ | $r_{young}$ | $MAE_{old}$ | $MAE_{young}$ | $RMSE_{old}$ | $RMSE_{young}$ |
|---|---|---|---|---|---|---|---|
| | EFN | .22 | .1 | .53 | .69 | .68 | .82 |
| | PercMot | .19 | .12 | .54 | .71 | .68 | .91 |
| | Power | .12 | .02 | .56 | .74 | .71 | .96 |
| RSFC | | | | | | | |
| | EFN | -.13 | .01 | .62 | .74 | .79 | .97 |

| | | | | | | | |
|---|---|---|---|---|---|---|---|
| | PercMot | .15 | -.04 | .54 | .77 | .69 | .97 |
| | Power | .10 | -.07 | .55 | .73 | .68 | .94 |
| ReHo | | | | | | | |
| | EFN | .02 | .12 | .59 | .72 | .73 | .93 |
| | PercMot | -.18 | -.11 | .6 | .77 | .76 | .99 |
| | Power | -.04 | .1 | .62 | .73 | .76 | .94 |
| fALFF | | | | | | | |
| | EFN | .08 | .1 | .55 | .66 | .69 | .87 |
| | PercMot | .05 | .06 | .54 | .66 | .69 | .88 |
| | Power | .05 | .08 | .56 | .66 | .7 | .88 |

*Note.* See Table S1.

Table S7

*Results of Complementary Machine-Learning-Adjusted t-Tests of the Effects of Modality, Age, Network, and Task Demand Level.*

| Factor | Comparison | t-value |
|---|---|---|
| Age | Old vs. Young | .06 |
| Demand Level | High vs. Low | 1.39 |
| Modality | GMV vs. RSFC | 2.67* |
| | GMV vs. ReHo | 4.96** |
| | GMV vs. fALFF | .854 |
| | RSFC vs. ReHo | 5.06** |
| | RSFC vs. fALFF | 4.63** |

| | | |
|---|---|---|
| | fALFF vs. ReHo | 5.30** |
| Network | EFN vs. PercMot | .073 |
| | EFN vs. Power | 1.36 |
| | PercMot vs. Power | 2.05* |

*Note.* EFN = executive-function-related network, PercMot = perceptuo-motor-related network, Power = Power et al.'s (2011) graph of putative functional areas, GMV = gray-matter volume, RSFC = resting-state functional connectivity, ReHo = regional homogeneity, fALFF = fractional amplitude of low frequency fluctuations.

**significant at $p < .001$

*significant at $p < .05$
